# BBSome-deficient cells activate intraciliary CDC42 to trigger actin-dependent ciliary ectocytosis

**DOI:** 10.1101/2023.08.13.553141

**Authors:** Avishek Prasai, Olha Ivashchenko, Kristyna Maskova, Marketa Schmidt Cernohorska, Ondrej Stepanek, Martina Huranova

## Abstract

Bardet-Biedl syndrome (BBS) is a pleiotropic ciliopathy caused by dysfunction of the BBSome, a cargo adaptor essential for export of transmembrane receptors from cilia. Although actin-dependent ectocytosis has been proposed as compensatory cellular process for defective cargo retrieval, the underlying molecular mechanism is poorly understood, particularly in the context of BBS pathology. In this study, we investigated how actin polymerization and ectocytosis are regulated within the cilium. Our findings reveal that ciliary CDC42, a RHO-family GTPase triggers *in situ* actin polymerization, ciliary ectocytosis, and cilia shortening in BBSome-deficient cells. Furthermore, activation of the Sonic Hedgehog pathway further enhances CDC42 activity in BBSome-deficient cilia, but not in healthy cells. Inhibition of CDC42 decreased the frequency and duration of ciliary actin polymerization events and lead to the accumulation of the GPR161 receptor in bulges along the axoneme in BBSome-deficient cells during Sonic Hedgehog signaling. Overall, our study identifies CDC42 as a key trigger of ciliary ectocytosis. Moreover, the hyperactivated ciliary CDC42-actin-ectocytosis axis in BBSome-deficient cells results in cilia shortening and loss of ciliary material, potentially impacting disease severity.

## Introduction

Primary cilia are microtubule-based organelles protruding from the cell surface. Primary cilia, referred to as ‘cilia’ onwards, are rich in signaling receptors, which sense various extracellular stimuli. Particular mutations in genes critical for cilia function cause pleiotropic human diseases, collectively called ciliopathies (reviewed in [1]). Bardet-Biedl syndrome (BBS) is a multi-organ ciliopathy caused by dysfunction of the BBSome, an octameric cargo adaptor involved in export of specific G-protein coupled receptors from primary cilia [2–6]. BBS presents with developmental and functional anomalies in the retina, brain, kidney, liver, heart, and other organs [7]. However, the molecular mechanisms of how the BBSome deficiency leads to the particular pathological outcomes in BBS are still incompletely understood.

The dysfunction of the BBSome results in the accumulation of signaling receptors in the cilia. This triggers the alternative pathway to shut down the signaling via the release of the ciliary receptors in ciliary vesicles [4, 6, 8]. BBSome-deficient cells have usually shorter cilia than wild type (WT) cells [9–12], which could be a consequence of the continuous ectocytosis.

The removal of the ciliary membrane via ectocytosis regulates ciliary signaling also under physiological conditions and drives cilia disassembly prior mitosis [4, 13–17]. The ectosomes are formed at the ciliary tip and are released upon an actin polymerization event inside cilia [13–15]. In the cytoplasm, actin polymerization is controlled by the GTPases of the RHO family, which cycle between the GTP and GDP loaded state [18]. RHO GTPases are vital for diverse cellular processes, including cell polarization, migration, and division, regulating cytoskeletal dynamics, membrane remodeling, and signaling pathways. Several RHO family members (CDC42, RHOA, RHOC, RAC1), their regulators and effectors were detected in the proteomic analyses of the ciliary content [19, 20]. Whether and how the individual RHO GTPases regulate actin polymerization in the cilia and the ectocytosis has not been addressed.

Here, we report that CDC42, a key regulator of cell polarity and actin-based morphogenesis [21], has an additional role as an essential trigger of the intraciliary actin polymerization during ectocytosis in BBSome-deficient cells.

## Methods

### Antibodies, dyes and reagents

Mouse anti-acetylated tubulin (1:50) was kindly provided by Dr. Vladimir Varga (Institute of Molecular Genetics of the Czech Academy of Sciences), rabbit anti-GPR161 (1:200) was kindly provided by Dr. Saikat Mukhopadhyay (UT Southwestern Medical Center). Rabbit anti-ARL13B (17711-1-AP; 1:2000) was purchased from Proteintech.

Secondary antibodies were as follows: anti-mouse Alexa Fluor 488 (Invitrogen, A11001; 1:1000), anti-rabbit Alexa Fluor 488 (Invitrogen, A11008; 1:1000), anti-mouse Alexa Fluor 555 (Invitrogen, A21422; 1:1000), anti-rabbit Alexa Fluor 555 (Invitrogen, A21428; 1:1000), anti-mouse Alexa Fluor 647 (Invitrogen, A21235; 1:1000) and anti-rabbit Alexa Fluor 647 (Invitrogen, A21245; 1:1000).

Phalloidin Texas Red (T7471; 1:250) and ProLong™ Gold antifade reagent with 4′,6-diamidino-2-phenylindole (DAPI) (P36941) or without (P36930) was purchased from Invitrogen.

### Cell cultures and treatments

Immortalized human retinal pigment epithelium cell line, hTERT-RPE1 (RPE1) (ATCC, CRL-4000) was kindly provided by Dr. Vladimir Varga (Institute of Molecular Genetics of the Czech Academy of Sciences). *BBS4* ^KO/KO^, *BBS1* ^KO/KO^ and *BBS7* ^KO/KO^ RPE1 cell lines were established previously [11]. Phoenix Eco cells were kindly provided by Dr. Tomas Brdicka (Institute of Molecular Genetics of the Czech Academy of Sciences). Mouse embryonic fibroblast (MEFs) lines were established previously [22]. All cell lines were cultured in complete Dulbecco’s modified Eagle’s medium (DMEM, Sigma, D6429 - 500 mL) supplemented with 10% fetal bovine serum (FBS), 100 U/mL penicillin (BB Pharma), 100 μg/mL streptomycin (Sigma-Aldrich), and 40 μg/mL gentamicin (Sandoz).

Smoothened agonist - SAG (566660; Sigma-Aldrich) was used at a concentration of 200 nM. CDC42 inhibitor – ML141 (4266; Tocris Biosciences) was used at 50 µM, ROCK-1 inhibitor - Y27632 (Y0503; Sigma-Aldrich) was used at 20 µM, RAC1 inhibitor (CAS1177865-17-6; Tocris Biosciences) was used at 100 µM, Cytochalasin D (C8273; Sigma-Aldrich) was used at 1 µM and ARP 2/3 complex inhibitor - CK666 (182515; Sigma-Aldrich) was used at 100 µM concentration. The treatments were for 2 h if not indicated otherwise.

### Cloning and gene transfections

*GPR161* ORFs was amplified from cDNA obtained from RPE1 cells, appended with mCherry coding sequence at the C terminus using recombinant PCR, and cloned into pMSCV-IRES-Thy 1.1 vector (Clonetech) using XhoI and ClaI restriction sites. CDC42 WT (12599) and DN (12601) plasmids were obtained from Addgene [23]. Ciliary targeting motif NPHP3 [1–203] [19] was fused to GFP-CDC42, WT and DN, N-terminally and subcloned into pMSCV-IRES-Thy 1.1 vector (Clontech) using EcoRI and ClaI restriction sites.

CDC42 Raichu probe was a kind gift from Prof. Michiyuki Matsuda (Kyoto University Graduate School of Medicine) [24]. Raichu probe was fused N-terminally with the ciliary targeting motif and sub-cloned into pMSCV-IRES-Thy 1.1 vector using EcoRI and ClaI restriction sites. CFP-PAK-CDC42 was amplified from the CDC42 Raichu probe and N-terminally tagged with the ciliary targeting motif and sub-cloned into pMSCV-IRES-Thy 1.1 using EcoRI and ClaI restriction sites.

LifeAct-TagRFP was a kind gift from Dr. Zdenek Hodny (Institute of Molecular Genetics of the Czech Academy of Sciences). mNeonGreen-ARL13B was a kind gift from Dr. Vladimir Varga (Institute of Molecular Genetics of the Czech Academy of Sciences).

The viral transduction of cell lines was done according to [25] with slight modifications. For viral particle production, Platinum Eco cells were seeded on 10 cm dish and allowed to reach confluency of 60-70%. 30 µg of plasmid DNA was transfected into cells using polyethyleneimine to generate retroviruses. Cells were incubated overnight and the media was changed to production media (DMEM+ATB+FBS) on the next day. MEFs were transduced with 2 mL supernatant containing the viral particles along with 8 µg/mL polybrene and analysed for transgene expression after 48 h. MEFs expressing CFP, YFP, mNeonGreen, mCherry, and/or TagRFP were bulk sorted using a FACSAria IIu (BD Biosciences).

### Immunofluorescence

RPE1 or MEF cells were seeded on 12-mm coverslips and serum starved for 24 h. After respective treatments, cells were fixed (4% formaldehyde) and permeabilized (0.2% Triton X-100) for 10 min. Blocking was done using 5% goat serum (Sigma, G6767-100 mL) in PBS for 15 min and incubated with primary antibody (1% goat serum/PBS) and secondary antibody (PBS) for 1 h and 45 min, respectively in a wet chamber. The cells were washed after each step in PBS three times. At last, the cells were washed in distilled H_2_O, air-dried, and mounted using ProLong™ Gold antifade reagent with DAPI (P36941; Invitrogen).

### Fluorescence microscopy

Image acquisition was performed on the Delta Vision Core microscope using the oil immersion objective (Plan-Apochromat 60× NA 1.42) and filters for DAPI (435/48), FITC (523/36), TRITC (576/89) and Cy5 (632/22). Z-stacks were acquired at 1024 × 1024-pixel format and Z-steps of 0.2 µm. Z-stacks were analysed using Fiji ImageJ software. Maximum intensity projections were used to quantify the frequency of ARL13B and GPR161 foci and GPR161 positive cilia and to measure the cilia length using the line tool in the Fiji ImageJ software.

### Expansion microscopy

Expansion microscopy of primary cilia was done as described previously [11]. RPE1 cells were cultured on 12 mm coverslips and serum starved for 24 h. Coverslips were fixed with 4% formaldehyde/4% acrylamide in PBS overnight and then washed 2× with PBS. The gelation was performed by incubating coverslips face down with 45 μL of monomer solution (19% (W/W) sodium acrylate, 10% (W/W) acrylamide, 0.1% (W/W) N,Ń-methylenbisacrylamide in PBS supplemented with 0.5% TEMED and 0.5% APS), in a pre-cooled humid chamber. After 1 min on ice, chamber was incubated at 37°C in the dark for 30 min. Samples in the gel were denatured in denaturation buffer (200 mM SDS, 200 mM NaCl, 50 mM Tris in ddH_2_O) at 95°C for 4 h. Gels were expanded in ddH_2_O for 1 h and then cut into 1×1 cm pieces. Pieces of gel were incubated with primary antibodies diluted in 2% BSA in PBS overnight at RT. After staining, shrunk pieces of gel were incubated in ddH_2_O for 1 h. After expansion, pieces of gel were incubated with secondary antibodies diluted in 2% BSA in PBS for 3 h at RT. Last expansion in ddH_2_O with exchange every 20 min was for 1 h until pieces of gel reached full size. Samples were imaged in 35 mm glass bottom dishes (CellVis) pre-coated with poly-L-lysine. During imaging, gels were covered with ddH_2_O to prevent shrinking. Expanded cells were imaged by confocal microscopy on Leica TCS SP8 using a 63× 1.4 NA oil objective with closed pinhole to 0.4 AU. Cilia images were acquired in Z-stacks at 0.1 μm stack size with pixel size 30-50 nm according to the cilia length. Images were de-convolved using Huygens Professional v. 21.04 software (Scientific Volume Imaging, Hilversum, Netherlands).

### Live cell imaging, data processing, and analysis

MEF cells expressing GPR161-mCherry or mNG-ARL13B and LifeAct-TagRFP were cultured on glass bottom dish (CellVis) until confluent. Cells were serum starved for 24 h for the induction of ciliogenesis. Before imaging, cells were treated with SAG and DMSO or CDC42 inhibitor, ML141 in FluoroBrite DMEM (Gibco, A1896701) and imaged immediately using Delta Vision Core microscope at 37°C with 5% CO_2_. Images were acquired using the oil immersion objective (Plan-Apochromat ×60, NA 1.42) in 1024 × 1024–pixel format and Z-steps of 0.2 μm. 7-8 imaging positions containing cilia were set. Imaging was performed in 2 min intervals to acquire a time-lapse video.

Imaging for GPR161-mCherry expressing cells was performed using the filter for mCherry (632/60) for 100 min (Figure 4C). Imaging for mNG-ARL13B and LifeAct-TagRFP expressing cells was performed using filters for FITC (523/36) and TRITC (576/89) for 120 min (Figure 5A, S5 and S6).

Time-lapse videos were de-convolved using Huygens Professional v. 21.04 (Scientific Volume Imaging, Hilversum, Netherlands) using the classic maximum likelihood estimation (CMLE). Signal to noise ratio was set to 40 with the area radius of 0.5 µm. One brick mode was used for the de-convolution and a maximum iteration of 40. De-convolved time-lapse videos were further analysed using Fiji ImageJ. For 3D analysis and visualization Imaris viewer 9.9.1 (Bitplane, Oxford Instruments plc) was used. Movies were extracted from de-convolved time-lapse videos. Time lapse video was cropped for the region of interest and time and animated with a rotation of 360°.

### Fluorescence lifetime imaging microscopy, data processing, and analysis

MEFs were seeded on 12-mm coverslips and serum starved for 24 h. To activate SHH pathway, cells were incubated with the 200 nM SAG for 2 h. The coverslips were fixed with 4% formaldehyde for 5 min, rinsed in PBS and water, air-dried and embedded in ProLong™ Gold antifade reagent and imaged on the same day. The FLIM-FRET measurements were done using Leica Stellaris 8 Falcon confocal microscope equipped with an oil immersion objective HC Plan-Apochromat 63× NA 1.4 oil, CS2 at room temperature. The acquisition and subsequent analysis was done using the built-in Phasor FLIM software tool [26]. Donor CFP fluorescence was excited by a pulsed (40MHz) white laser tuned at 440 nm, and emitted photons between 457 nm and 488 nm were collected (max. 500 photons per pixel) using the HyD X2 detector in photon counting mode. The acquisition was performed in 512×512-pixel format with pinhole 2.5 AU, at a speed of 400 Hz in bidirectional mode and 8-bits resolution. In each experiment, non-transfected cells were used to estimate the autofluorescence and the N-CFP-PAK-CDC42 expressing WT MEFs were used to estimate the lifetime of donor CFP only. These two parameters were used to estimate the FRET fraction and lifetime of the donor CFP (Raichu probe) in the phasor space [26].

### Statistical analysis

Statistical analysis was performed using GraphPad Prism Version 5.04. All statistical tests were two-tailed. The statistical tests are indicated in the respective Figure legends.

## Results

### Differential roles of RHO family members in the regulation of the cilia length

The BBSome functions mostly as a retrograde cargo adaptor for the G-protein coupled receptors (GPCRs) in cilia [3]. Loss of BBSome alters the ciliary export leading to ectocytosis of accumulated cargoes [4]. As actin polymerization is typically triggered by different members of RHO family GTPases in different contexts, we aimed to identify the RHO family member(s) triggering the actin polymerization during ciliary ectocytosis.

We hypothesized that the cilia shortening in the BBSome-deficient cell lines, previously observed by others [9, 10, 12] and us [11] (Fig. 1A and B) is caused by continuous ectocytosis. This was supported by the presence of buldge-like structures, possibly nascent ectosomes, in the cilia of *BBS4* ^KO/KO^ and *BBS1* ^KO/KO^ RPE1 cell lines (Fig. 1C, Fig. S1A). To identify the putative RHO GTPase responsible for the cilia shortening in BBSome-deficient cells by ectocytosis, we treated the cells with well-established inhibitors of the three major RHO GTPases: Y27632 (inhibits ROCK1, a downstream effector of RHOA [9]), ML141 (inhibits CDC42 [27]) and CAS 1177865-17-6 (shortly CAS; inhibits RAC1 [28]) (Fig. 1D, Fig. S1B). The inhibition of ROCK1 prolonged cilia both in WT, *BBS1*^KO/KO^, and *BBS4*^KO/KO^ RPE1 cells (Fig. 1D, E) [9], consistent with the role of RHOA-dependent cortical F-actin network in the regulation of the cilia length [29, 30]. Accordingly, the treatment with Cytochalasin D and ARP2/3 inhibitor resulted in cilia prolongation in both cell lines (Fig. S1B-C). In contrast, RAC inhibition induced only very subtle changes in cilia length (Fig. S1D). Only the inhibition of CDC42 prolonged cilia specifically in the BBSome-deficient cells, whereas the treatment had no effect in WT cells (Fig. 1D, F). Moreover, we observed a higher frequency of foci with accumulated ciliary membrane marker, ARL13B at the ciliary tips of *BBS4* ^KO/KO^ cells compared to the WT cells (Fig. 1G-H, Fig. S1E). Inhibition of CDC42 increased the amount of the ARL13B foci specifically in the *BBS4* ^KO/KO^ cells (Fig. 1H).

**Figure 1.**
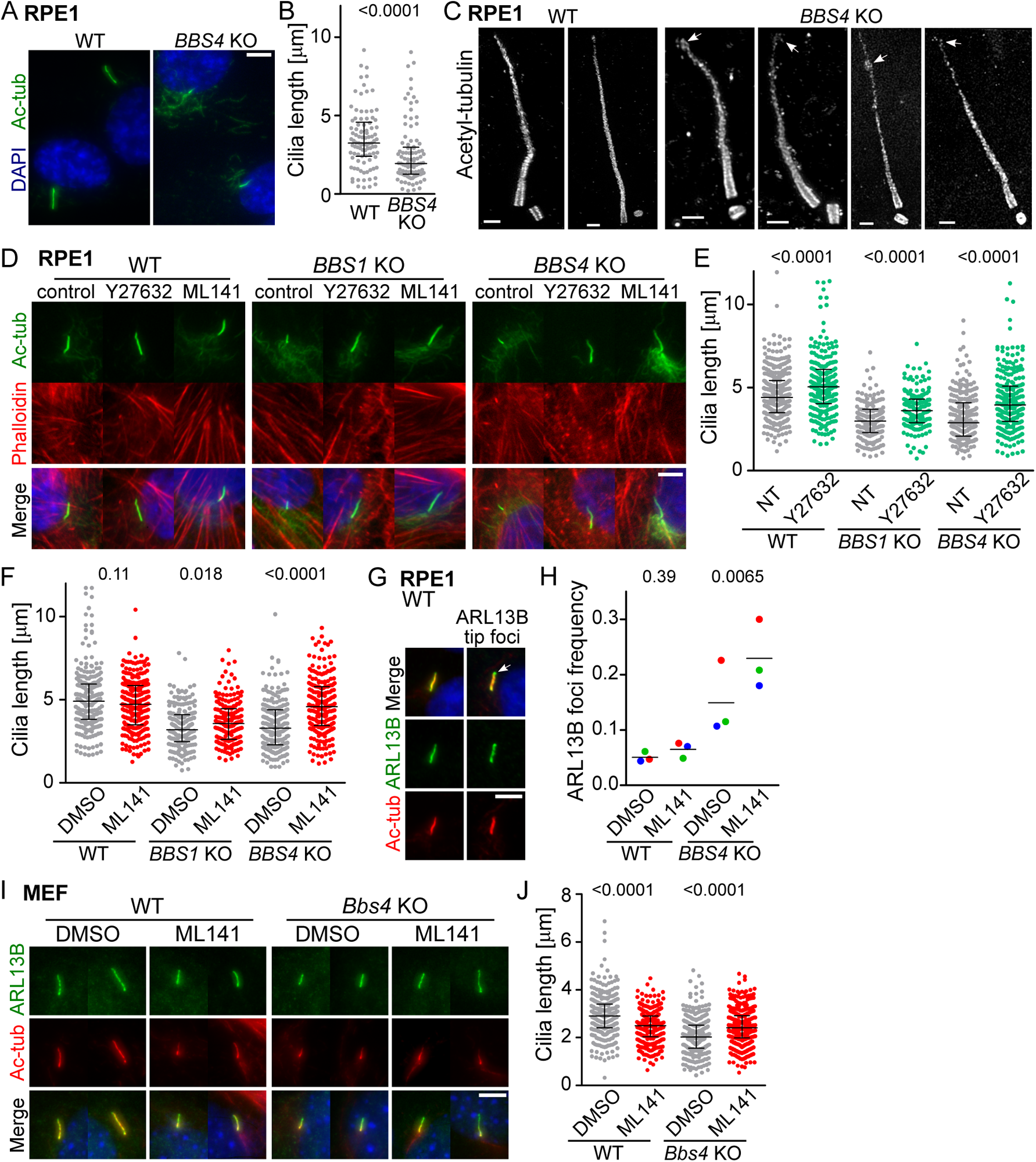
RHO family members regulate differentially the cilia length. (A) Representative micrographs and (B) quantification of the cilia length, stained with acetylated tubulin (Ac-tub), of the WT and *BBS4*^KO/KO^ RPE1 cells. Scale bar, 5 µm. Medians with interquartile range from three independent experiment (n = 96 cilia). (B) Expansion microscopy of the cilia of the WT and *BBS4*^KO/KO^ RPE1 cells stained for acetylated tubulin marking the axoneme. White arrows show ciliary bulges observed in *BBS4*^KO/KO^ cells. Scale bar, 2 µm. (C) Representative micrographs of cilia (Ac-tub) and F-actin (Phalloidin) in WT, *BBS1*^KO/KO^ and *BBS4*^KO/KO^ RPE1 cells treated with the ROCK1 inhibitor Y27632 or CDC42 inhibitor ML141. Scale bar, 5 µm. (D) Quantification of the cilia length in the WT, *BBS1*^KO/KO^ and *BBS4*^KO/KO^ RPE1 cells non-treated (NT) or treated with Y27632. Medians with interquartile range from three independent experiments (n = 160-280 cilia). (E) Quantification of the cilia length in the WT, *BBS1*^KO/KO^ and *BBS4*^KO/KO^ RPE1 cells treated with vehicle or ML141. Medians with interquartile range from three independent experiments (n = 180-280 cilia). (F) Micrographs depicting cilium (Ac-tub) without and with the ARL13B positive foci at the tip (arrow) in WT RPE1 cells. Scale bar, 5 µm. (G) Quantification of the frequency of ciliary tip foci observed in G. Mean of three independent experiments (n = 100-200 cilia). (H) Representative micrographs and (J) quantification of the cilia length, in WT and *Bbs4*^KO/KO^ MEFs treated with vehicle or ML141 stained for acetylated tubulin (Ac-tub) and ARL13B. Scale bar, 5 µm. Medians with interquartile range from three independent experiments (n = 300-400 cilia). Statistical significance was calculated using the two-tailed Mann-Whitney test (B, E, F, J) or two-tailed paired t-test (H). Merged micrographs show nuclei staining by DAPI – blue (D, G, I).

In the next step, we utilized the WT and *Bbs4*^KO/KO^ mouse embryonic fibroblasts (MEFs) [22] to verify our observations in a primary cell line model. In WT MEFs, inhibition of CDC42 resulted in cilia shortening (Fig. 1I, J), indicating that ciliogenesis in MEFs is dependent on the CDC42-exocyst mediated delivery of ciliary vesicles to cilia as proposed previously [31]. Yet, inhibition of CDC42 increased the cilia length in *Bbs4*^KO/KO^ MEFs corroborating the unprecedented additional function of CDC42 in ectocytosis (Fig. 1I, J).

Overall, these findings imply that the loss of BBSome triggers CDC42-mediated ectocytosis and concurrent shortening of cilia in various cell lines.

### CDC42 controls GPR161 and cilia dynamics in BBSome-deficient cells

Sonic Hedgehog (SHH) signaling initiates the BBSome-dependent removal of GPR161 from the cilia, which triggers the downstream signaling [5]. In line with the previous results [4, 6], we observed ciliary accumulation of GPR161 in *Bbs4*^KO/KO^ MEF cells in the steady state, which was not altered after activation of the SHH pathway via the SMO agonist (SAG) (Fig. 2A-B). We frequently observed GPR161 localized to specific foci at the ciliary tip of SAG-stimulated *Bbs4*^KO/KO^ MEFs (Fig. 2C), indicating ongoing ectocytosis. SHH signaling decreased the cilia length in WT and *Bbs4*^KO/KO^ MEFs (Fig. 2A, D-E). However, the inhibition of CDC42 by ML141 prevented the SAG-induced cilia shortening in *Bbs4*^KO/KO^ MEFs, but not in WT cells (Fig. 2D-E).

**Figure 2.**
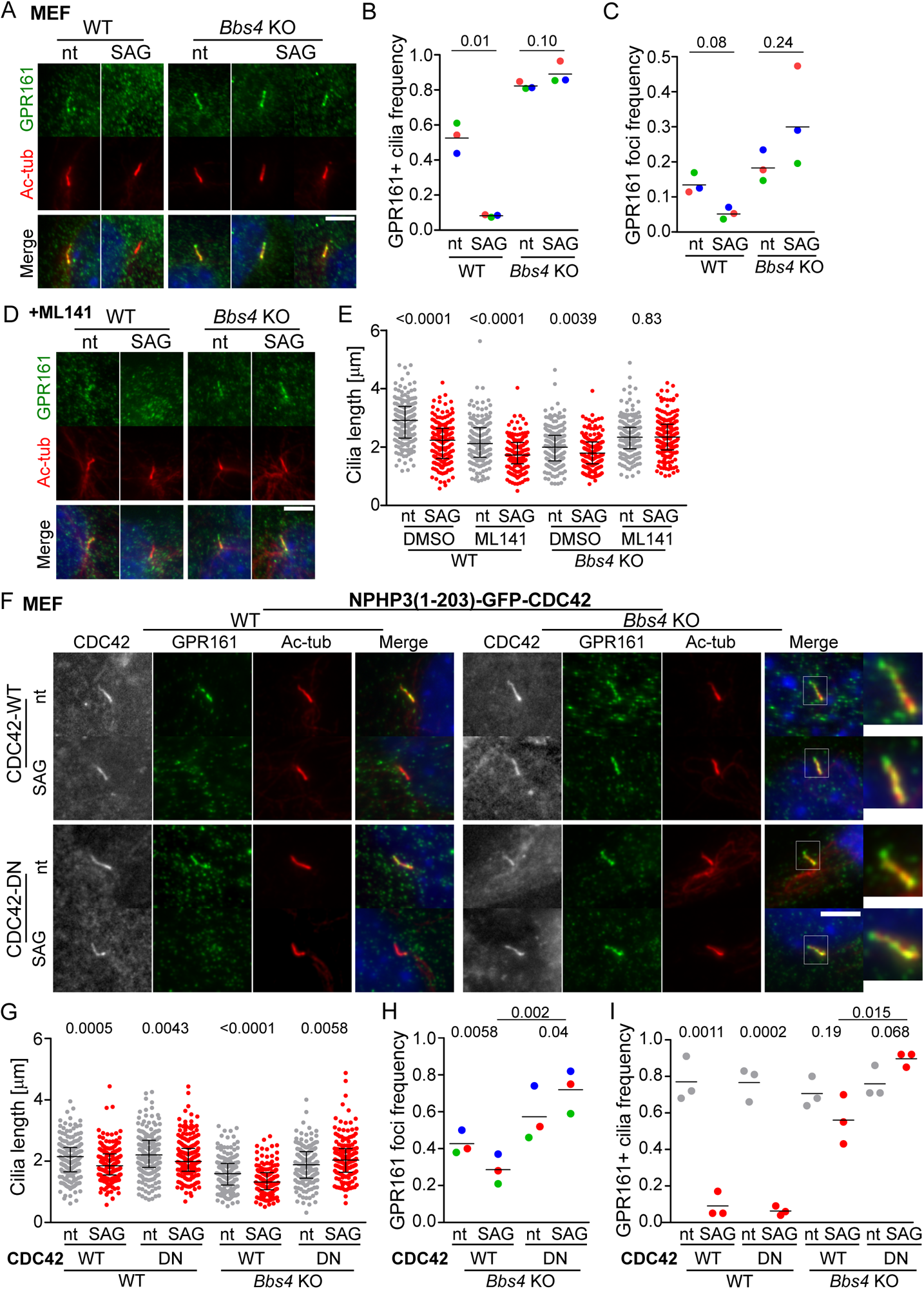
CDC42 controls GPR161 and cilia dynamics in BBSome-deficient cells. (A) Representative micrographs show cilia (Ac-tub) and GPR161 in non-treated (nt) and SAG induced WT and *Bbs4*^KO/KO^ MEFs. Scale bar, 5 µm. (B) Quantification of the frequency of GPR161 positive cilia and (C) of the frequency of GPR161 foci at the cilia tip in non-treated (nt) and SAG induced WT and *Bbs4*^KO/KO^ MEFs in A. Mean of three independent experiments (n = 160-210 cilia). (C) Representative micrographs show cilia (Ac-tub) and GPR161 in non-treated (nt) and SAG induced WT and *Bbs4*^KO/KO^ MEFs concomitantly treated with ML141. Scale bar, 5 µm. (D) Quantification of the cilia length, in non-treated (nt) and SAG induced WT and *Bbs4*^KO/KO^ MEFs concomitantly treated with vehicle or ML141. Medians with interquartile range from three independent experiments (n = 170-190 cilia). (E) Representative micrographs show cilia (Ac-tub), and localization of GPR161 (Merge and insets), in non-treated (nt) and SAG induced WT and *Bbs4*^KO/KO^ MEFs expressing the cilia targeted CDC42 WT or DN variant. Scale bar, 5 µm. (F) Quantification of the cilia length, in non-treated (nt) and SAG induced WT and *Bbs4*^KO/KO^ MEFs expressing cilia targeted CDC42 WT or DN variant. Medians with interquartile range from three independent experiments (n = 160-220 cilia). (G) Quantification of the frequency of GPR161 tip foci in non-treated (nt) and SAG induced *Bbs4*^KO/KO^ MEFs expressing cilia targeted CDC42 WT or DN variant. Mean of three independent experiments (n = 160-190 cilia). (H) Quantification of the frequency of GPR161 positive cilia in non-treated (nt) and SAG induced WT and *Bbs4*^KO/KO^ MEFs expressing cilia targeted CDC42 WT or DN variant. Mean of three independent experiments (n = 160-190 cilia). Statistical significance was calculated using two-tailed paired t-test (A-C, H-I) or two-tailed Mann-Whitney test (D, G). Merged micrographs show nuclei staining by DAPI - blue (A, D, F).

To address whether cytoplasmic or ciliary CDC42 promotes the cilia shortening in BBSome-deficient cells, we expressed WT or dominant negative (DN) CDC42 with a ciliary targeting motif [19] in the WT and *Bbs4* deficient MEFs (Fig. 2F). We observed that CDC42-DN abrogated cilia shortening and promoted the formation of the GPR161 foci in the cells lacking BBSome in the steady state (Fig. 2F-H). Activation of the SHH pathway lead to comparable cilia shortening and GPR161 removal in CDC42-WT and DN expressing WT MEFs (Fig. 2G, I). In contrast, CDC42-DN blocked SAG-induced cilia shortening and removal of ciliary GPR161 in the *Bbs4*^KO/KO^ MEFs, (Fig. 2G-I).

Altogether, these data show that ciliary CDC42 promotes ectocytosis leading to the cilia shortening in cells lacking the BBSome in the steady state and during SHH signaling.

### CDC42 is hyperactivated in cilia in BBSome deficient cells during SHH signaling

To directly assess the activity of CDC42 inside the cilia, we expressed the Raichu-CDC42 FRET-FLIM genetic probe [24] appended with an N-terminal ciliary anchor [19] (N-Raichu-CDC42) in WT and *Bbs4*^KO/KO^ MEFs (Fig. 3A-B). Activation of CDC42 in the probe leads to an intramolecular interaction with PAK resulting in the Förster resonance energy transfer (FRET) from a donor (CFP) to acceptor (YFP) and thus decrease in donor lifetime. We measured the activity of CDC42 in cilia in non-stimulated cells and after SHH activation (Fig. 3C-D, S2A). The estimated FRET efficiency corresponded to the fraction of the reporter CDC42 molecules in the active conformation (Fig. 3D). We detected basal activity of CDC42 in cilia in both the WT and *Bbs4*^KO/KO^ cells as depicted by the general decrease in CFP lifetime when compared to the no-FRET reference control N-CFP-PAK-CDC42 (Fig. 3D, S2A). In the steady state, the CDC42-GTP fraction was slightly higher in the *Bbs4*^KO/KO^ cells (Fig. 3D), which could explain the CDC42 dependent cilia shortening in these cells. The activity of CDC42 was unaffected by the SHH signaling in the WT cells (Fig. 3D), indicating that the concomitant cilia shortening is CDC42 independent (Fig. 2F). On the other hand, activation of the SHH pathway in *Bbs4*^KO/KO^ cells lead to substantial increase in CDC42 activity (Fig. 3D). These data document that the BBSome deficiency induces the CDC42 activity in cilia in the steady state and particularly during ciliary signaling.

**Figure 3.**
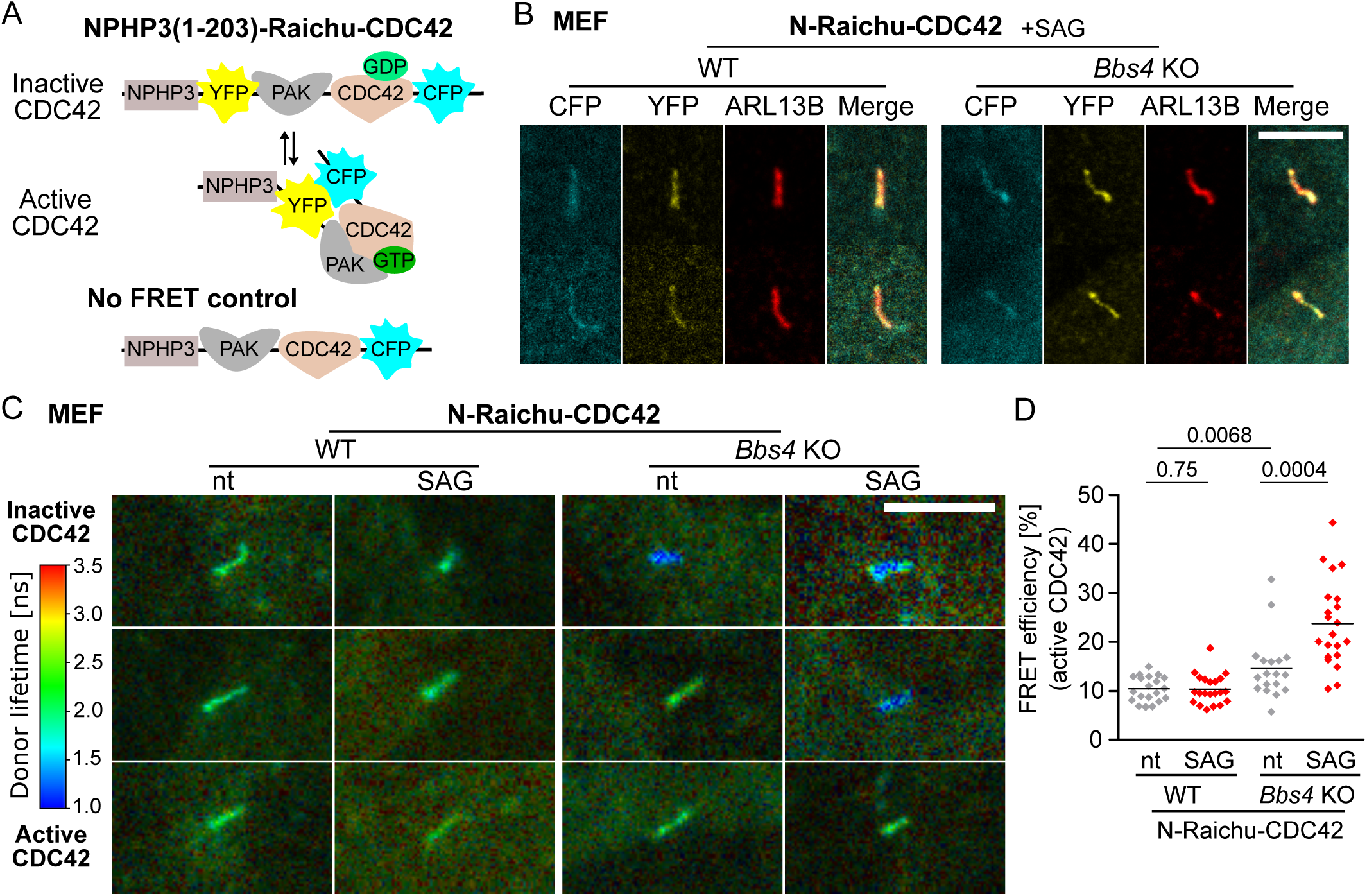
CDC42 is hyperactivated in cilia in BBSome-deficient cells. (A) Schematic representation of the Raichu-CDC42 FRET probe with the ciliary targeting motif NPHP3 (1-203) and the no-FRET control lacking YFP. Activation of CDC42 is measured as decrease in FRET donor (CFP) lifetime. (B) Representaive micrographs show cilia (ARL13B) and the cellular localization of N-Raichu-CDC42 probe (YFP and CFP) in WT and *Bbs4*^KO/KO^ MEFs induced with SAG. Scale bar, 5μm. (C) Representative micrographs show donor lifetime values in non-treated (nt) and SAG induced WT and *Bbs4*^KO/KO^ MEFs expressing N-Raichu-CDC42 determined by the FLIM-FRET analysis in fixed cells. Lifetimes are shown in pseudocolours ranging from blue to red. (D) The FRET efficiency measured in cilia in non-treated (nt) and SAG induced WT and *Bbs4*^KO/KO^ MEFs expressing N-Raichu-CDC42. Mean of three (WT) and four (KO) independent experiments (n = 18-21 cilia). Statistical significance was calculated using the two-tailed Mann-Whitney test.

### SHH signaling induces CDC42-dependent cilia shortening in BBSome-deficient cells

To explore whether the excess of the BBSome-dependent cargoes in cilia triggers the CDC42 mediated ectocytosis, we overexpressed GPR161-mCherry in *Bbs4*^KO/KO^ MEFs (Fig. 4A). We observed that the GPR161-mCherry accumulated at the ciliary tips in the steady state and that the frequency of these GPR161-mCherry positive foci increased upon SHH induction (Fig. 4A-B). Inhibition of CDC42 under both conditions further increased the frequency of the GPR161 positive bulges (Fig. 4A-B). Notably, we observed that the overexpressed GPR161 accumulated in bulges also along the axoneme when CDC42 was inhibited over the course of SHH signaling (Fig. 4A).

**Figure 4.**
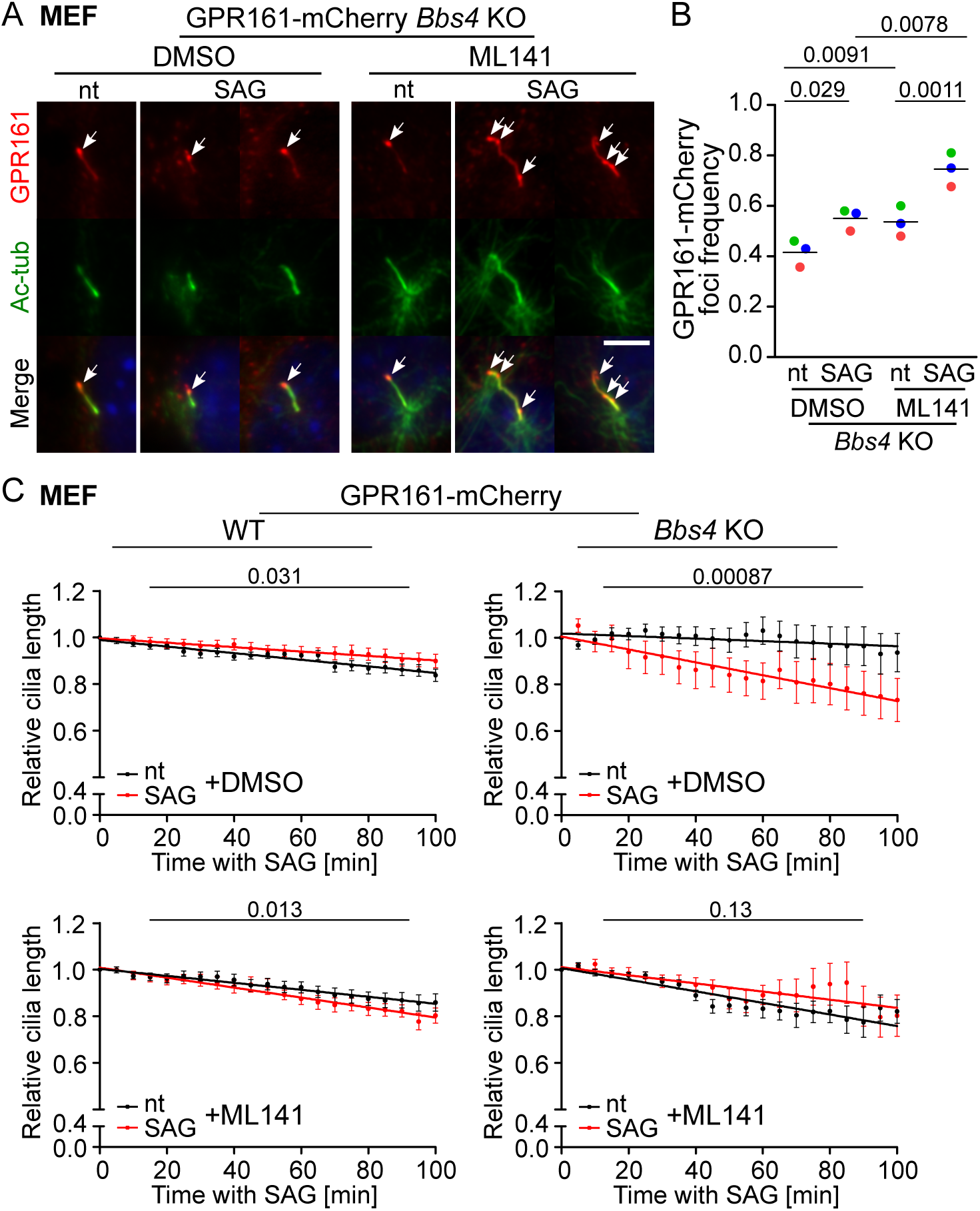
SHH signaling induces CDC42 dependent cilia shortening in BBSome-deficient cells. (A) Representative micrographs and (B) quantification of the frequency of GPR161-mCherry foci (arrows) in cilia (Ac-tub) in non-treated (nt) and SAG induced *Bbs4*^KO/KO^ MEFs concomitantly treated with vehicle or ML141. Scale bar, 5 μm. Mean of three independent experiments (n = 100-160 cilia). Statistical significance was calculated using the two-tailed paired t-test. (B) Plots depict the average dynamics of the cilia length in non-treated (nt) and SAG induced WT and *Bbs4*^KO/KO^ MEFs treated concomitantly with vehicle or ML141 and imaged for 100 min. The length of the cilium was normalized to the length measured at time 0 min. In total 17-30 cilia were monitored per condition in two independent experiments. The data points were fitted using the linear regression and the statistical significance of the slope of the regression lines was calculated for the indicated conditions.

We monitored the cilia length during the course of SHH activation in cells pre-treated with the CDC42 inhibitor via time-lapse live cell imaging (Fig. 4C, Fig. S2B). We observed that activation of the SHH pathway triggered cilia shortening in the *Bbs4*^KO/KO^, but not in the WT MEFs, and that this cilia shortening was blocked by the inhibition of CDC42 (Fig. 4C, Fig. S2B-C). Altogether, these data show that CDC42 controls the signal dependent shortening of primary cilia via ectocytosis in the BBSome deficient cells.

### CDC42 is required for actin polymerization inside the cilia

Our data indicated that the accumulation of signaling receptors activate CDC42 to trigger the ectocytosis. In the next step, we examined whether CDC42 induces actin polymerization inside the cilia. We expressed the membrane marker ARL13B-mNeonGreen (ARL13B-NG) and actin binding protein LifeAct-TagRFP in the WT and *Bbs4*^KO/KO^ MEFs. We monitored the dynamics of the ciliary membrane and actin polymerization upon SHH activation in the absence or presence of ML141 using time-lapse imaging (Fig. 5A-B, Fig. S3, S4 and MOV. 1-4). In several cases, we observed actin polymerization with following ectocytosis, which we defined as visible separation of the ARL13B positive membrane segments (Fig. 5B, MOV. 1-2). We detected more actin polymerization events in the cilia of the *Bbs4*^KO/KO^ cells in comparison to WT cells (Fig. 5C, Fig. S3A, S4A). Interestingly, the overall dynamics and duration of the F-actin patches was very variable (Fig. 5D). We observed that over the course of SHH activation, the inhibition of CDC42 reduces the frequency of the ciliary actin polymerization events (Fig. 5C, Fig. S3B, S4B, and MOV. 3-4) and their duration (Fig. 5D).

**Figure 5.**
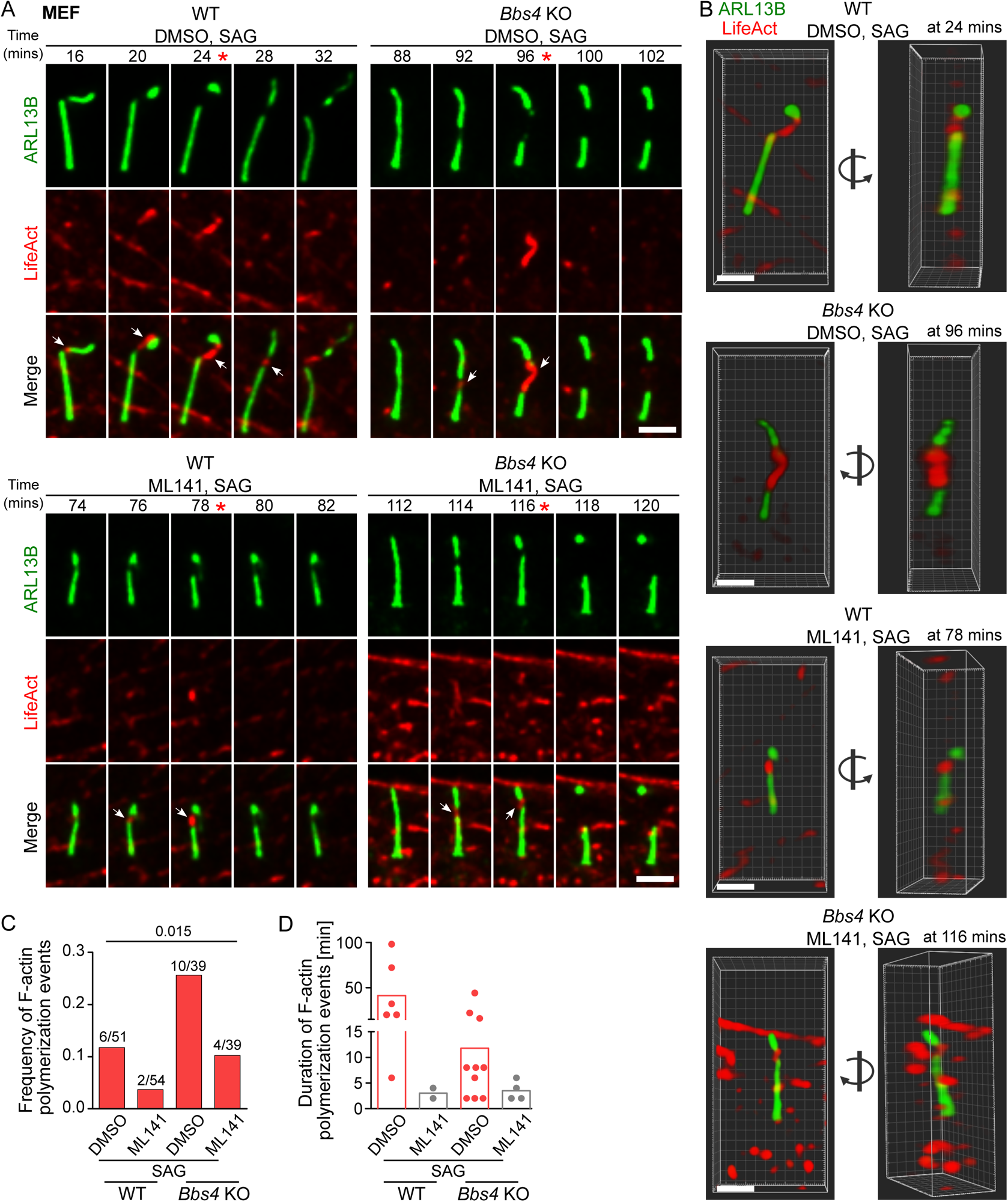
CDC42 is required for actin polymerization inside the cilia. (A) Representative micrographs depicting ciliary ectocytosis and actin polymerization detected by live cell imaging of WT and *Bbs4*^KO/KO^ MEFs expressing mNG-ARL13B and LifeAct-TagRFP. The cells were treated with SAG and vehicle (top) or ML141 (bottom) and imaged every 2 min for 2 h. White arrows point to the F-actin polymerization events. Red asterisks indicate the frames used for 3D visualization in B. Scale bar, 2 μm. (B) 3D visualization of the F-actin patches and ciliary membrane in the WT and *Bbs4*^KO/KO^ cells at the indicated time points of the live cell imaging in A. Left micrographs show the front view of the cilium and right micrographs show a 90° rotated view in the direction indicated by the arrows. Scale bar, 2 μm. (C) The graph shows the fraction of cilia with actin polymerization events observed in WT and *Bbs4*^KO/KO^ MEFs expressing mNG-ARL13B and LifeAct-TagRFP upon treatment with SAG and vehicle or ML141 during the live cell imaging in A. The number of observed actin polymerization events out of the total count of imaged cilia from four independent experiments is shown. Statistical analysis was done using the contingency table and Chi-square test. (D) The plot depicts the duration of the observed actin polymerization events detected in cilia in the WT and *Bbs4*^KO/KO^ MEFs expressing mNG-ARL13B and LifeAct-TagRFP upon treatment with SAG and vehicle or ML141 during the live cell imaging in A.

Altogether, our study revealed that the CDC42 triggers actin polymerization inside the cilia to promote ectocytosis in the BBSome-deficient cells in the steady state and during SHH signaling.

## Discussion

In this study, we have shown that the inhibition of CDC42 prevented cilia shortening specifically in the BBSome-deficient cells. The concomitant inhibition of CDC42 and activation of the SHH pathway blocked the signal-induced ectocytosis of GPR161 and led to its accumulation in bulges along the axoneme. Using cilia-targeted dominant negative CDC42 mutant, we revealed that intraciliary CDC42 triggers ciliary ectocytosis that controls the overall ciliogenesis process in the BBSome-deficient cells. In some instances, we observed growing F-actin patches prior to ectocytosis. This suggests that actin microfilaments could exert pushing forces to release the ectosomes from cilia [32, 33]. Accordingly, the actin polymerization events were rare and only short-lived upon the CDC42 inhibition, indicating that CDC42 induces and sustains ciliary F-actin structures within cilia to trigger ectocytosis. Overall, we showed that the loss of the retrograde cargo adaptor BBSome leads to the hyperactivation of CDC42 within cilia, both in the steady state and during the SHH signaling. Hyperactive CDC42 triggers excessive F-actin mediated ectocytosis, which leads to cilia shortening, the typical phenotype linked to BBS [9–12].

CDC42 along with other actin regulators was detected in the ciliary proteome [19, 20]. The localization of CDC42 within cilia could be mediated by the Par6-dependent translocation from the basal body [34]. Par6, together with aPKC, links CDC42 activity also to the non-canonical WNT/PCP signaling, which facilitates polarized reorganization of the microtubules [35]. It remains to be elucidated whether the hyperactivation of ciliary CDC42 is a consequence of persistent signaling of ciliary GPCRs in the BBSome-deficient and/or it is directly linked to the BBSome deficiency via its involvement in the WNT/PCP pathway [36, 37].

Although studies on actin polymerization in the cilia are only emerging [13, 15, 38, 39], roles of cortical F-actin and its regulators from the RHO family in ciliogenesis are well established [9, 16, 29–31, 40–42]. While RHOA-mediated actin depolymerisation promotes centrosome polarization, cilia assembly and elongation [29, 30, 40], polymerization and branching of actin is associated with cilia disassembly and inhibited ciliogenesis [41, 43]. This is in line with the previous and our observations that the inhibition of ROCK1, a downstream kinase of RHOA, prolongs cilia length in both WT and BBSome-deficient cells [9].

CDC42 is so far the only GTPase with documented localization at the basal body [42]. CDC42 controls multiple aspects of ciliogenesis including its involvement in vesicular transport [31], endocytosis [41] and signal transduction [42]. However, the outcomes of perturbing CDC42 function have been contradictory [31, 42]. These discrepancies could be potentially explained by the intricate interplay between the extraciliary and intraciliary functions of CDC42 in ciliogenesis with variable outcomes, depending on the specific context and cell type. CDC42 knockdown or expression of a non-targeted DN CDC42 mutant impairs the CDC42 activity both in the cilia and in the cell body and thus, cannot discriminate between the specific and perhaps counteracting roles of these two pools [31, 42]. In contrast, our approach with ciliary-targeted DN CDC42 addresses exclusively the function of CDC42 inside the cilia.

Although the molecular etiology of particular BBS symptoms in tissues is largely unexplained, it is plausibly linked to defects in specific signaling pathways [44]. The BBSome deficiency presents with short cilia and mislocalization of the BBSome dependent cargoes, mostly G-protein coupled receptors (GPCRs). In cells lacking the BBSome, several GPCRs such as GPR161, GPR19, and D1 accumulate in the cilia [6, 45, 46], which corresponds to the well-described role of the BBSome in the retrograde transport [3, 4, 8, 47]. However, some other receptors disappear from cilia in the BBSome-deficient cells and tissues, such as SSTR3, MCHR1, and NPY2 in neurons of mouse models of BBS [46, 48, 49]. This was originally explained by the proposed role of the BBSome in the import of these receptors into the cilia [2]. However, the BBSome-mediated ciliary cargo import has never been clearly documented on a molecular level. It is thus possible that the absence of these receptors from the cilia can be caused by the BBSome-deficiency indirectly via the enhanced ectocytosis which might remove these receptors in a by-stander manner. In this scenario, the CDC42-mediated ectocytosis in BBSome-deficient cells would be a pathological mechanism connected with the BBS pathology.

BBSome-deficient photoreceptors accumulate mislocalized proteins in their outer segments, which is a highly specialized cilium [50–52]. As the formation of the photoreceptor discs resembles modified ectocytosis [53], it is possible that the BBSome deficiency interferes with this process which leads to the dysfunction of photoreceptors and eventualy causes retinal dystrophy, one of the hallmarks of the BBS [51, 54].

The BBSome-deficiency leads to morphological changes also in motile cilia in brain ependymal layer and in the airways [55–57], which are responsible for the flow of cerebrospinal fluid and the transport of mucus and protection from respiratory infections, respectively. The motile cilia of several examined *Bbs* KO mice formed bulges at the ciliary tip filled with vesicles, which was accompanied by altered cilia beating [55, 57]. It is possible that the CDC42-mediated ectocytosis or a related process are triggered in the motile cilia upon the BBSome dysfunction.

Altogether, enhanced and altered ectocytosis in BBSome-deficient cells could contribute to the manifestation of BBS symptoms. Thus, the inhibition of the increased ectocytosis could be a novel potential therapeutic strategy in the BBS.

## Acknowledgement

This project has received funding from the Czech Science Foundation (21-21612S to MH), core funding provided by the Institute of Molecular Genetics of the Czech Academy of Sciences (RVO 68378050), project National Institute of Virology and Bacteriology (Programme EXCELES, LX22NPO5103 to OS) - funded by the European Union - Next Generation EU, Charles University Grant Agency (386321 to AP and OI) and EU Horizon 2020 Research and Innovation programme (MSCF 847693 to MSC). We acknowledge the Light Microscopy Core Facility, IMG, Prague, Czech Republic, supported by MEYS – LM2023050 and RVO – 68378050-KAV-NPUI, for their support with the widefield and confocal imaging and data analysis presented herein.

## Author contribution

MH conceived the study and was in charge of the overall direction and planning. AP, OI and MH performed the basic immunofluorescence, live cell imaging, FLIM microscopy and data analysis, MSC performed the expansion microscopy experiments, MH and AP performed cloning and together with KM and OI generated the cell lines present in this study. AP, OS and MH wrote the manuscript with the contribution of all the other authors.

