## Supplemental information for "BBSome-deficient cells activate intraciliary CDC42 to trigger actin-dependent ciliary ectocytosis"

**Supplemental Figure 1. Actin regulators regulate the cilia length in different manner**

(A) Expansion microscopy of the cilia of the *BBS1*$^{KO/KO}$ and *BBS7*$^{KO/KO}$ RPE1 cells stained for the acetylated tubulin marking the axoneme. White arrows point to bulges at the cilia tips. Scale bar, 2 µm.

(B) Representative micrographs of the cilia (Ac-tub) and actin cytoskeleton (Phalloidin) of the WT RPE1 cells treated with inhibitors of ROCK1/RHOA (Y27632), CDC42 (ML141), RAC1 (CAS1177865-17-6 (CAS)), ARP2/3 (CK666) and actin polymerization (Cytochalasin D). Scale bar, 5 µm.

(C) Quantification of the cilia length in the WT and *BBS4*$^{KO/KO}$ RPE1 cells treated with vehicle or CK666 or Cytochalasin D (CytoD). Medians with interquartile range from three independent experiments (n = 220-320 cilia).

(D) Quantification of the cilia length in the WT and *BBS4*$^{KO/KO}$ RPE1 cells treated with vehicle or CAS. Medians with interquartile range from three independent experiments (n = 250-280 cilia).

(E) Representative micrographs of the cilia (Ac-tub, ARL13B) of the WT and *BBS4*$^{KO/KO}$ RPE1 cells treated with vehicle or ML141. Scale bar, 5 µm.

Statistical significance was calculated using the two-tailed Mann-Whitney test (C, D). Merged micrographs show nuclei staining by DAPI – blue (E).

**Supplemental Figure 2. CDC42 hyperactivation drives cilia shortening in BBSome deficient cells upon SHH triggering**

(A) The donor lifetime values and measured in cilia in non-treated (nt) and SAG induced WT and *Bbs4*$^{KO/KO}$ MEFs expressing N-Raichu-CDC42 and values measured for the no-FRET control expressed in WT MEFs. Mean of three (WT) and four (KO) independent experiments (n = 18-21 cilia).

(B) Representative micrographs of the cilia in non-treated (nt) and SAG induced WT and *Bbs4*$^{KO/KO}$ MEFs expressing GPR161-mCherry, concomitantly treated with vehicle or ML141 and imaged every 2 min for 100 min. Frames were extracted from the live cell videos (17-30 cilia per condition in two independent experiments). Scale bar, 2 µm.

(C) Plots depict the variable dynamics of the length of the individual cilia in non-treated (nt) and SAG induced WT and *Bbs4*$^{KO/KO}$ MEFs expressing GPR161-mCherry, concomitantly treated with vehicle or ML141 and imaged every 2 min for 100 min in B. The length of the cilium was normalized to the length measured at time 0 min. In total 17-30 cilia were monitored per condition in two independent experiments.

**Supplemental Figure 3. CDC42 is required for actin polymerization in cilia in WT cells.**

(A) Representative micrographs show F-actin polymerization events observed in cilia in WT MEFs expressing mNeonGreen-ARL13B (green) and LifeAct-TagRFP (red) treated with SAG and vehicle. Frames were extracted from time-lapse videos (51 in total, 6 actin polymerization events, four independent experiments). Scale bar, 2 μm.

(B) Representative micrographs show F-actin polymerization events observed in WT MEFs expressing mNeonGreen-ARL13B (green) and LifeAct-TagRFP (red) treated with SAG and ML141. Frames were extracted from time-lapse videos (54 in total, 2 actin polymerization events, four independent experiments). Scale bar, 2 μm.

Maximum intensity projections of the z-stacks were done using Fiji ImageJ software and the intensities for both channels were adjusted post acquisition for better visualization.

**Supplemental Figure 4. CDC42 is required for actin polymerization in cilia in BBSome deficient cells.**

(A) Representative micrographs show F-actin polymerization events observed in *Bbs4*$^{KO/KO}$ MEFs expressing mNeonGreen-ARL13B (green) and LifeAct-TagRFP (red) treated with SAG and vehicle. Frames were extracted from time-lapse videos (39 in total, 10 actin polymerization events, four independent experiments). Scale bar, 2 μm.

(B) Representative micrographs show F-actin polymerization events observed in *Bbs4*$^{KO/KO}$ MEFs expressing mNeonGreen-ARL13B (green) and LifeAct-TagRFP (red) treated with SAG and ML141. Frames were extracted from time-lapse videos (39 in total, 4 actin polymerization events, four independent experiments). Scale bar, 2 μm.

Maximum intensity projections of the z-stacks were done using Fiji ImageJ software and the intensities for both channels were adjusted post acquisition for better visualization.

**Movie 1.**

3D visulization of the actin polymerization event in the cilium of WT MEFs expressing mNeonGreen-ARL13B (green) and LifeAct-TagRFP (red) treated with SAG and vehicle. Time lapse videos were cropped for region of interest and time and rotation of 360° vertically is shown. Playback speed is 24 frames per second. Scale bar, 1 μm.

**Movie 2.**

3D visulization of actin polymerization event in the cilium of *Bbs4*$^{KO/KO}$ MEFs expressing mNeonGreen-ARL13B (green) and LifeAct-TagRFP (red) treated with SAG and vehicle. Time lapse videos were cropped for region of interest and time and rotation of 360° vertically is shown. Playback speed is 24 frames per second. Scale bar, 1 μm.

**Movie 3.**

3D visulization of actin polymerization event in the cilium of WT MEFs expressing mNeonGreen-ARL13B (green) and LifeAct-TagRFP (red) treated with SAG and ML141. Time lapse videos were cropped for region of interest and time and rotation of 360° vertically is shown. Playback speed is 24 frames per second. Scale bar, 1 μm.

**Movie 4.**

3D visulization of actin polymerization event in the cilium of *Bbs4*$^{KO/KO}$ MEF cells expressing mNeonGreen-ARL13B (green) and LifeAct-TagRFP (red) treated with SAG and ML141. Time lapse videos were cropped for region of interest and time and rotation of 360° vertically is shown. Playback speed is 24 frames per second. Scale bar, 1 μm.

Supplemental Figure 1

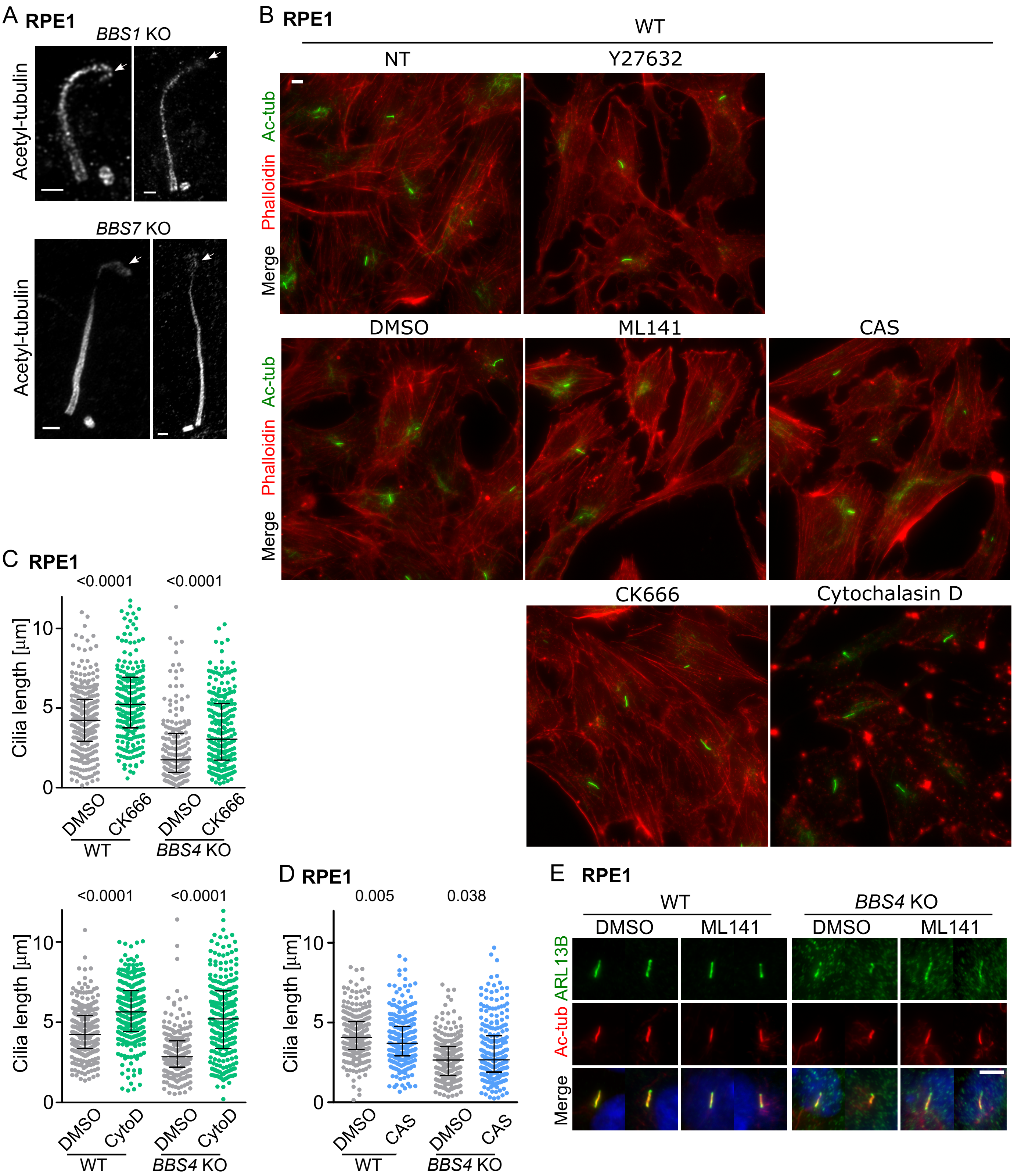

Supplemental Figure 2

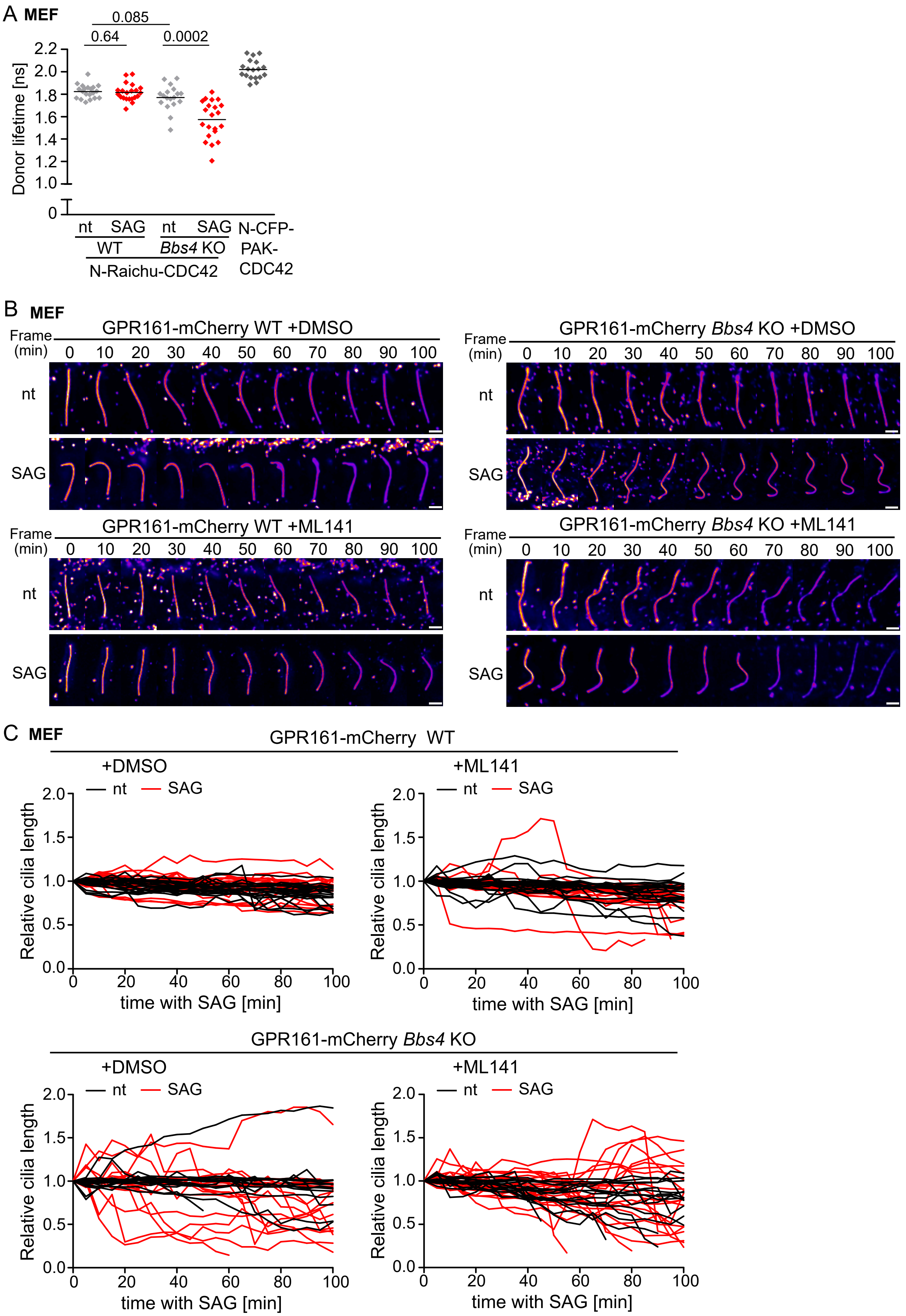

Supplemental Figure 3

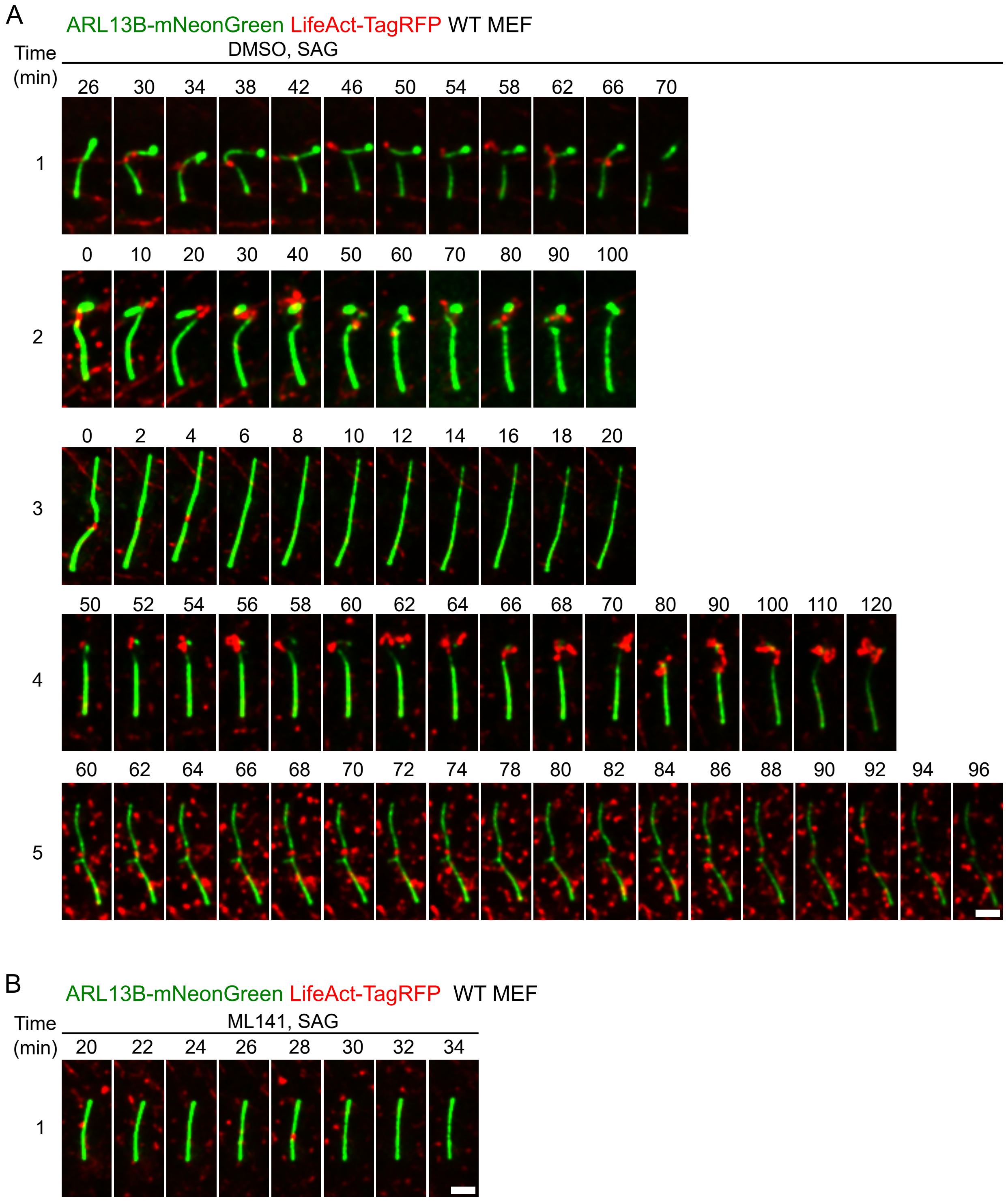

Supplemental Figure 4

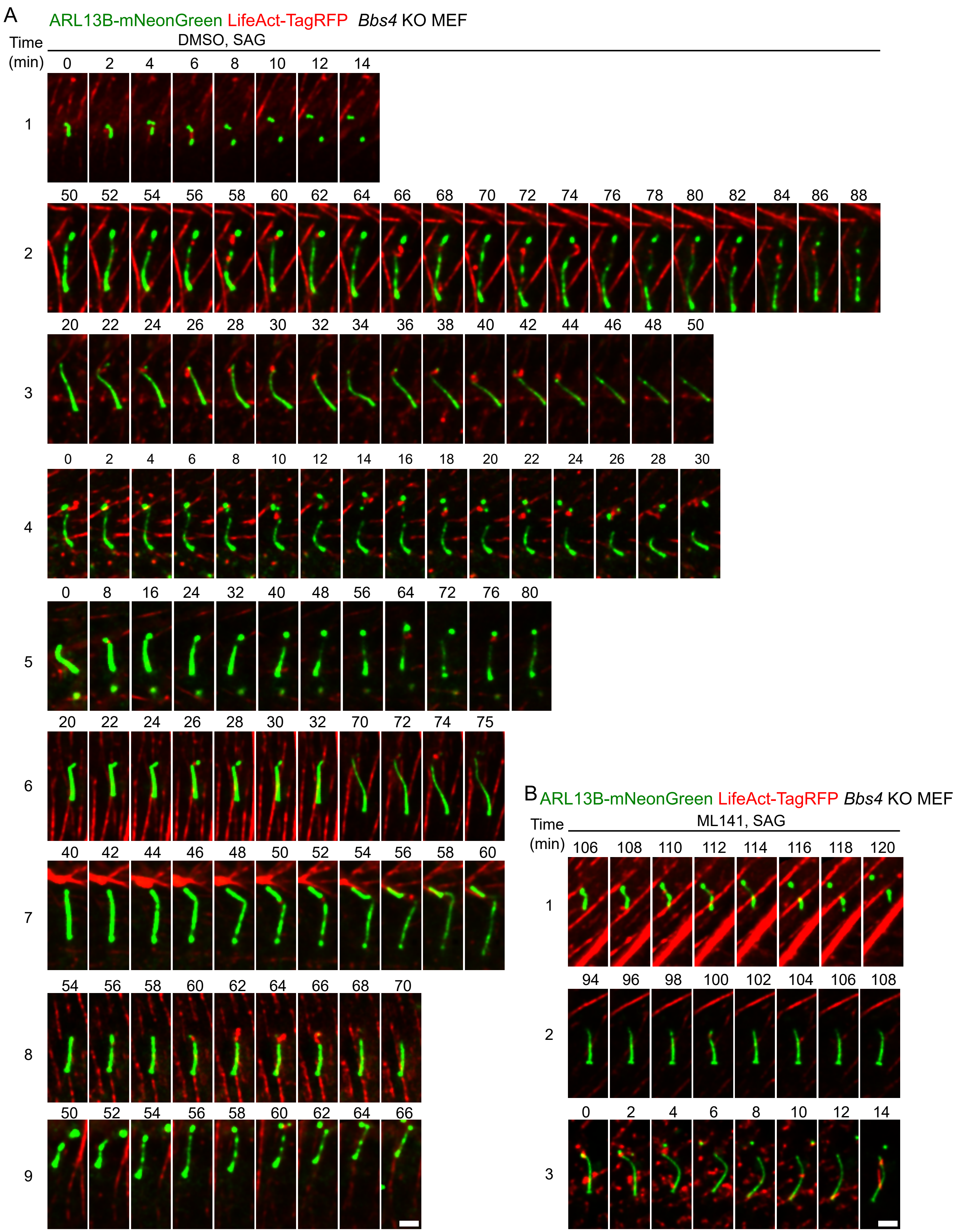
